# Spatial transcriptomics of tumor microenvironment in formalin-fixed paraffin-embedded breast cancer

**DOI:** 10.1101/2020.01.31.928143

**Authors:** Lou Romanens, Prasad Chaskar, Jean-Christophe Tille, Stephan Ryser, Nicolas Liaudet, Ketty Hu-Heimgartner, Killian Heimgartner, Gurkan Kaya, Petros Tsantoulis, S. Intidhar Labidi-Galy

## Abstract

Tumor samples are conserved in clinical practice in formalin-fixed paraffin-embedded (FFPE) blocks. Formalin fixation chemically alters nucleic acids, rendering transcriptomic analysis challenging. RNA-sequencing is usually performed on tumor bulk, without distinction of cell subtypes or location. Here we describe the development of a robust method for RNA extraction and exome-capture RNA-sequencing of laser-capture microdissected tumor cells (TC) and stromal immune cells (TIL) based on their morphology. We applied this method on 7 tumor samples (surgical or core needle biopsy) of triple-negative breast cancer (TNBC) stored in FFPE blocks over 3-10 years. Unsupervised clustering and principal component analysis showed a clear separation between gene-expression profile of TIL and TC. TIL were enriched in markers of B cells (*CD79B, PAX5 and BLNK*) and T cells (*CD2, CD3D and CD8B*) whereas tumor cells expressed epithelial markers (*EPCAM, MUC1* and *KRT8*). Microenvironment cell populations-counter (MCP)-counter deconvolution showed an enrichment in adaptive immune cell signatures in microdissected TIL. Transcripts of immune checkpoints were differentially expressed in TIL and TC. We further validated our results by qRT-PCR and multispectral immunohistochemistry. In conclusion, we showed that combining laser-capture microdissection and RNA-sequencing on archived FFPE blocks is feasible and allows spatial transcriptional characterization of tumor microenvironment.

## Introduction

Solid tumors are composed of cancer cells and non-cancer cells such as immune cells, fibroblasts and endothelial cells (1, 2) that vary in proportions and state from one tumor to another. These cells compose the tumor microenvironment (TME) and can have a tumor-promoting or tumor-suppressive role (3). For instance, stromal tumor-infiltrating lymphocytes (TIL) are associated with good prognosis in patients with triple-negative breast cancers (TNBC) and predict response to chemotherapy (4) whereas cancer-associated fibroblasts have a deleterious effect (5).

Bulk RNA-sequencing is an accessible method for studying tumor transcriptome. Although it might be possible to algorithmically deconvolute the abundance of TME components from bulk data (6, 7), this analysis cannot resolve fine details such as the activation state of immune cells and is challenging to validate experimentally. In addition, the spatial relationship of tumor components, for instance tumor-infiltrating versus tumor-excluded lymphocytes, is obvious under microscopy, but cannot be inferred from bulk data. Single-cell RNA sequencing is a powerful method to investigate cell-to-cell variability and discover biologically relevant cell subtypes but it has been limited so far to the analysis of fresh/frozen tissues and does not permit spatial characterization of TME. Importantly, single-cell RNA-sequencing cannot easily recapitulate the complexity of an heterogeneous tumor from a few thousand cells and has not been proven to accurately capture cell proportions, possibly overestimating the abundance of some cell types such as immune cells (8).

Laser capture microdissection (LCM) is a technique that allows to collect separately distinct cell populations under direct microscopic visualization. One of the advantages of LCM is that specific cell subtypes can be recognized based on their morphology on hematoxylin or immunohistochemically-stained tissue sections (9). The selected cells are cut from tissue section using a laser beam and collected separately from the rest of the section. This approach provides a pool of cells enriched for a cell subtype of interest, from which RNA can be extracted for sequencing. An important advantage of LCM is that it enables the reuse of clinical material, which is usually formalin-fixed paraffin-embedded (FFPE) for diagnostic purposes. Formalin fixation allows better preservation of tissue morphology but leads to cross-linking and fragmentation of nucleic acids, making downstream sequencing more challenging (10). To overcome the challenges of working with RNA from FFPE samples, we optimized multiple steps of LCM and RNA extraction processes. We demonstrated the feasibility of performing RNA-sequencing from laser-capture microdissected stromal TIL and tumor cells from FFPE samples of TNBC (Figure 1). Here, we performed an exome-capture RNA-sequencing method that captures the coding regions of fragmented RNA. We validated this approach by qRT-PCR on a selected list of genes and by multispectral immunohistochemistry. Overall, we propose a method that renders feasible spatial transcriptomic analyses of TME in FFPE tumor samples.

**Figure 1:**
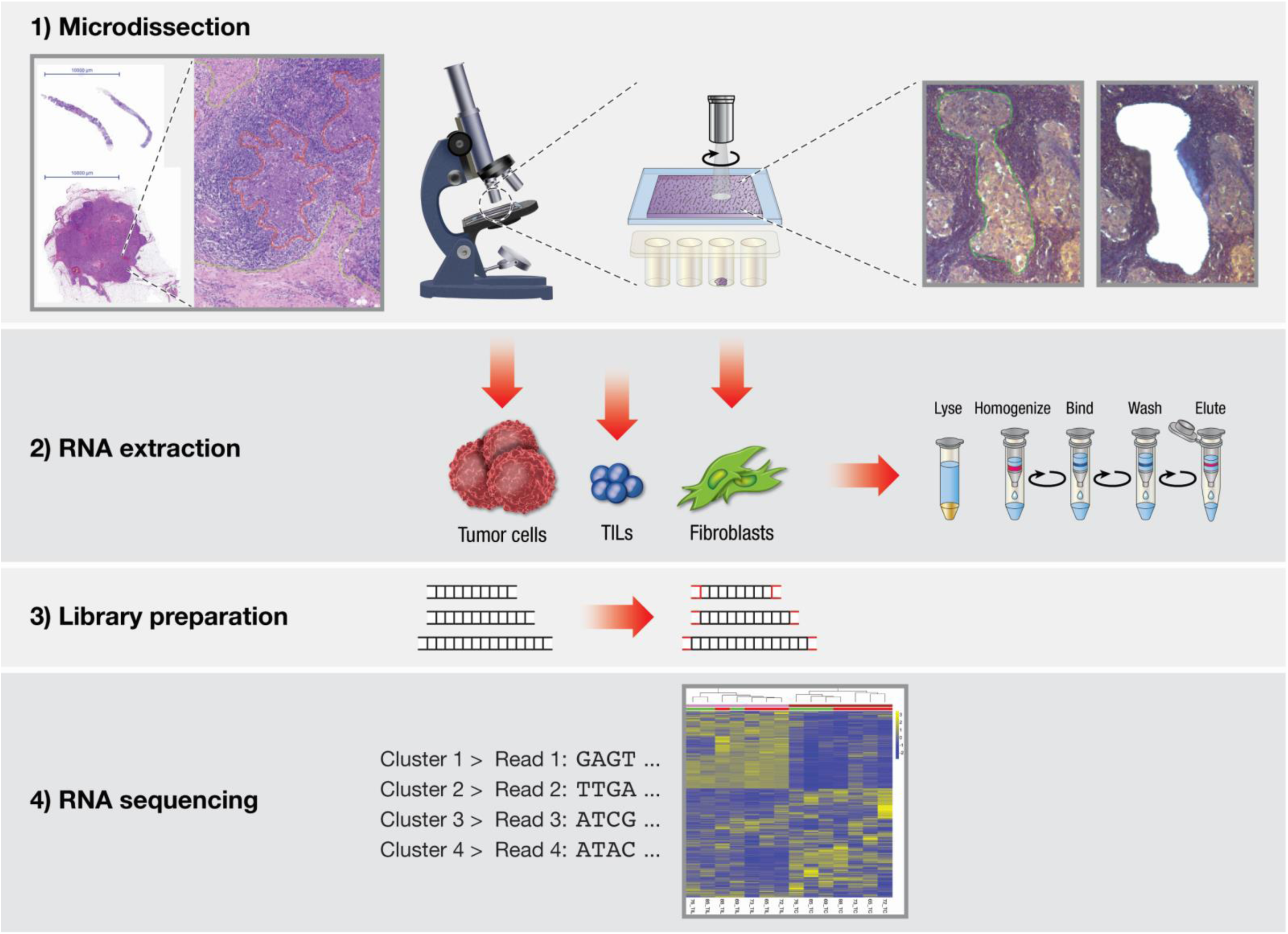
Workflow of laser capture microdissection and RNA sequencing. Step 1: TNBC tissue sections were cut, mounted on a polyethylene terephthalate membranes (PET) slide and stained with hematoxylin followed by laser-capture microdissection (LCM) procedure that includes manual delimitation of TC, TIL and Fib under direct microscopic visualization. Cells of interest were selected using a drawing tool on the Leica Laser Microdissection software. Selected regions of interest were cut from tumor sections with a laser beam and collected within the cap of a PCR tube placed under the PET slide. Step 2: total RNA from microdissected TC, TIL and Fib is extracted. Step 3: library preparation included the conversion of RNA into cDNA fragments and the addition of adaptors. Step 4: libraries are subjected to sequencing. TILs = tumor-infiltrating lymphocytes.

**Figure 2:**
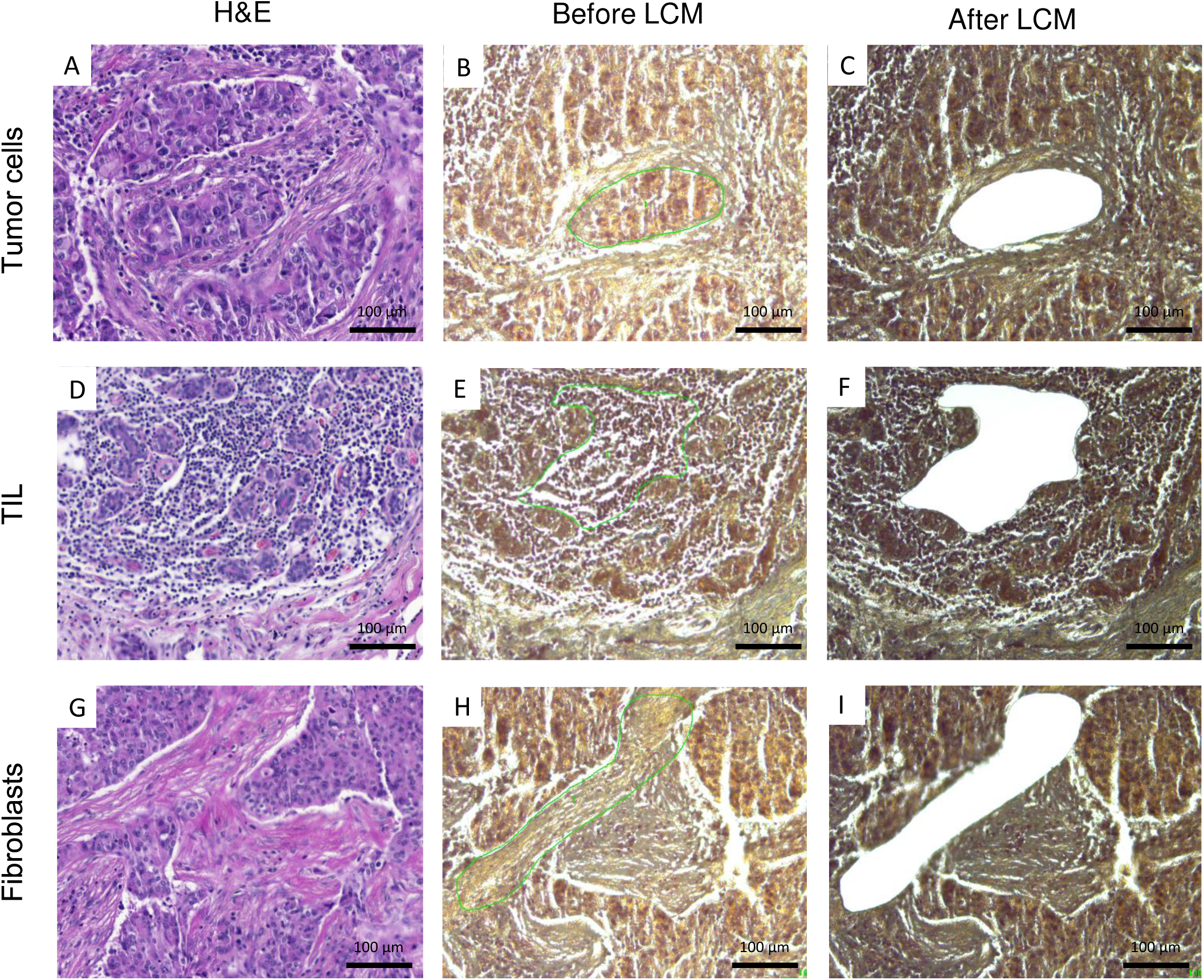
Laser-capture microdissection of different cell subpopulations based on their morphology. **A, D&G.** H&E-stained TNBC section containing the cells to be microdissected (upper panels: tumor cells; middle-panels: tumor-infiltrating lymphocytes; lower panels: fibroblasts). **B, E&H.** Selection of the region containing the cells of interest (green line) for laser-capture microdissection. **C, F&I.** Visualization of the tissue post-microdissection.

## Results

### Optimization of laser capture microdissection

Formalin fixation induces chemical cross-linking and RNA fragmentation, rendering subsequent genomic and transcriptomic analysis of FFPE material challenging. Thus, we optimized the tissue preparation process before LCM in order to maximize RNA yield while preserving RNA quality. The standard protocol for hematoxylin and eosin (H&E) staining used in clinical pathology lasts 27 min. However, prolonged immersion of tissue sections in clearing agent such as xylene further degrades nucleic acids (11). Therefore, we tested three different dyes (hematoxylin, eosin and crystal violet) and varying durations of incubation (supplementary Table 1 **and** supplementary Figure 1A). We aimed to shorten the duration of staining procedure while conserving maximal contrast in order to distinguish cell subtypes based on their morphology. Adding crystal violet dye or Scott’s tap water to the staining process led to low contrast images. The optimal staining was obtained with protocol b (supplementary Table 1) that had shorter duration of staining (15 min) and renders good contrast (supplementary Figure 1A).

Second, we determined the maximal tissue section thickness that allows the collection of a maximum number of cells and adequate visualization of cells’ morphology. Tissue sections of 6 and 8 µm thickness showed similar cell morphologies compared to the standard H&E tissue section of 3 μm whereas tissue section of 10 µm had too many “compacted” cells causing reduced contrast between tumor cells (TC), cancer-associated fibroblasts (Fib) and tumor-infiltrating lymphocytes (TIL) (supplementary Figure 1B). Therefore, we chose 8 µm as the optimal thickness for subsequent experiments.

Third, we aimed to determine the optimal surface of area to microdissect. A typical human cell contains 10-30 pg of RNA (12). However, the number of cells per surface varies according to tissue density. We estimated the number of cells to 7’000 per mm^2^ in TNBC sample. Accordingly, the quantity of RNA that would be theorically extracted was estimated to vary between 70 and 210 ng/mm^2^. We set up the minimal surface to be microdissected to 2.5 mm^2^ from 8 µm thick tissue sections of a one-year-old FFPE block which corresponds to an expected amount of 172-500 ng of RNA (Supplementary Figure 2).

### Optimization of RNA extraction

We sought to optimize the RNA extraction procedure from microdissected area of 2.5 mm^2^ with the objective of collecting RNA of the best possible quality and quantity. Minimal requirements for exome-capture RNA-sequencing with the TruSeq® protocol are: a) RNA integrity number (RIN) > 1.4, b) the percentage of fragments over 200 nucleotides (DV_200_) > 30% and c) a minimum yield of 40 ng depending on RNA quality (13). Following the RNeasy FFPE Qiagen® kit’s protocol (supplementary Figure 3A), RNA yield was undetectable using an elution volume of 30 µl for a tissue section surface of 2.5 mm^2^. Thus, we tested several modifications of the protocol in order to optimize RNA yield and quality. First, the eluted volume was reduced to 15 µl and mechanical digestion by stirring (400 rpm) was added, and subsequently concentrated using a vaccum concentrator centrifuge to a final volume of 4 µl. This rendered RNA detectable (yield 18-40 ng). Second, we increased the duration of enzymatic digestion. The original RNeasy protocol recommends tissue digestion for 15 min at 56 °C with proteinase K. We tested four different durations of digestion with proteinase K (15 min, 1h, 3h and overnight). RNA yield tended to increase with prolonged digestion, whereas DV_200_ initially reached a maximum at 3h but declined after overnight digestion. RIN was not affected by the duration of tissue digestion (supplementary Figure 3B-D). Thus, 3h was set as the optimal duration of digestion to collect RNA with sufficient quality and quantity. These modifications led to the final protocol (supplementary Figure 3E) that was tested on seven FFPE samples of TNBC aged from 3 to 10 years. Sufficient RNA from microdissected TIL and TC was collected from either core needle biopsies (n=2) or surgical biopsies (n=5) for subsequent sequencing (Supplementary Table 2). Extracting RNA from microdissecting Fib was more challenging: sufficient RNA yield was collected from only two out of seven tumors (supplementary Figure 3F). The age of the block did not impact RNA yield or RIN (supplementary Figure 3G-H) but we noticed a decrease of DV_200_ in older blocks (supplementary Figure 3I). *Library preparation and exome-capture RNA sequencing*

The library was prepared with the TruSeq® protocol on 16 samples: 7 TIL, 7 TC and 2 Fib (Supplementary Table 2). Exome-capture RNA sequencing was performed in multiplex, yielding an average of 49.3 M paired-end stranded reads. All samples passed the per base and per sequence quality score criteria (mean Phred quality score≥39) of the FastQC tool (Supplementary Table 3). On average, 76% of reads successfully aligned with the reference transcriptome (GRCh38 cDNA) using Salmon.

### Exploratory data analysis

Transcript abundance was estimated with Salmon and summarized at the gene level using the “tximport” tool from the DEseq2 package. The Principal Component Analysis (PCA) clearly separated TC, TIL and Fib. Technical replicates (same sample run in two different lanes) were in good agreement (Supplementary Figure 4) and were subsequently merged for better visualization (Figure 3A). These results suggest that TC samples are more similar to each other than they are to TIL and Fib. The same holds true for TIL and Fib samples. The PCA showed that the two Fib samples were placed closed to the TIL on the primary PCA axis, indicating a higher similarity among the Fib and TIL at the gene expression level (Figure 3A). The two Fib samples were excluded from further transcriptomic analyses. The distribution of TC and TILs samples according to the PCA was in agreement with the dendrogram and the heatmap generated based on correlation distances between samples (Figure 3B).

**Figure 3:**
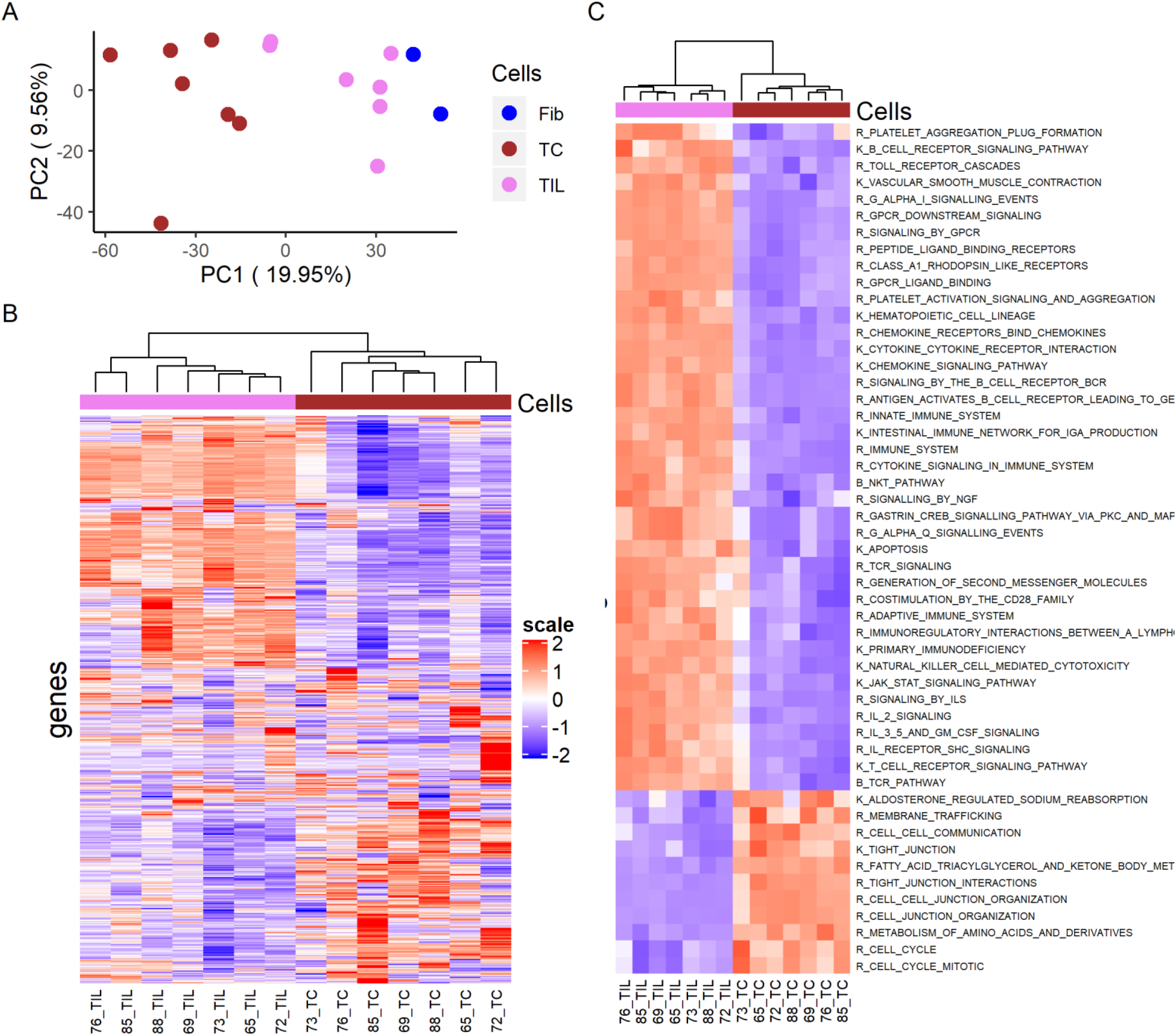
Differential gene expression in microdissected TC and TIL. **A.** Principal component analysis PCA of 16 microdissected samples based on the 500 most varying genes. **B.** Heatmap based on the hierarchical clustering of 7 TC (in brown) and 7 TIL (in pink) **C.** Heatmap based on Gene Set Variant Analysis, pathways enriched in TC and TIL. Color scale: red, high relative expression; blue, low relative expression. PC=Principal Component, R=Reactome, B=Bioconduct, K=KEGG. TC = tumor cells, TIL = tumor-infiltrating lymphocytes, Fib = fibroblasts.

### Transcriptomic characterization of TC and TIL

Pathway annotation using Gene set variant analysis (GSVA) estimates the enrichment of specific gene-sets on a sample-by-sample basis. Several canonical pathways related to immune cell functions such as TCR signaling, NKT signaling, BCR signaling, JAK-STAT signaling, IL-2/3/5 signaling were enriched in TIL whereas pathways related to cell cycle, cell junction organization, membrane trafficking were significantly enriched in TC (Figure 3C).

### Differential expression analysis

We performed differential expression analysis (DEA) comparing TC and TIL. In total, we identified 1776 genes that were differentially expressed between TC and TIL: 1003 genes were upregulated in TIL while 773 genes were upregulated in TC (**Supplementary Tables 4 and 5**). We observed that several genes such as *MUC1, TFAP2A, MAL2 and ELF3* already reported to be expressed in breast cancer (14–16) and particularly TNBC (17), were among the most significantly highly expressed genes in microdissected TC compared to TIL (**Supplementary table 4**). In addition, TC expressed several epithelial markers like *EPCAM, KRT8, KRT18* and *KRT19*. Among top upregulated genes in TIL, there were several genes known to be B cells-related (*CD79A, CD79B, PAX5, BLNK, and MS4A1*) and T cells-related genes (*CD2, CD3D, CD3E, CD3G, and CD8B*) (**Supplementary table 5**).

T cell co-signaling (co-stimulatory and co-inhibitory) receptors are cell-surface molecules that have a crucial role in regulating T cell function toward effector or suppression (18). We found that most of T cells co-stimulation markers such as *CD27*, *CD28* and *SLAMF1* and several co-inhibition markers/immune checkpoint inhibitors like *CTLA4* and *TIGIT* were substantially increased in microdissected TIL (Figure 4A and B) (19). PD-L1 *(CD274)* was numerically higher in TIL but the difference was not significant (*p*=0.39). Among immune checkpoint inhibitors, CD160 and BTLA are both ligands for the same receptor HVEM (*TNFRSF14*): we found that *BTLA* (*p*=1.36×10^-4^) was significantly upregulated in TIL but not *CD160* (Figure 4B), suggesting that within the context of TNBC BTLA could be more involved in tumor immunosuppression.

**Figure 4:**
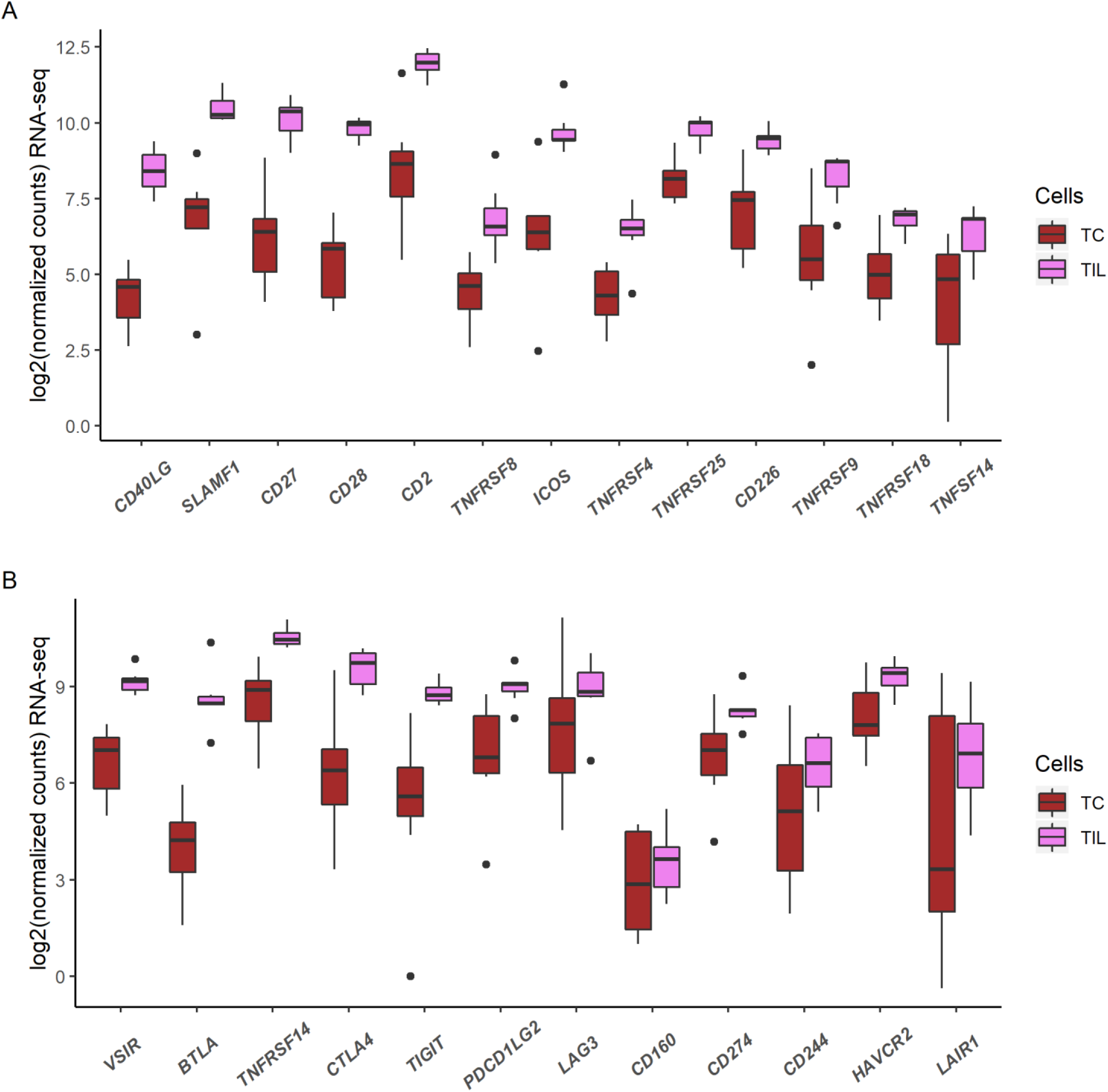
Expression level of T cells co-signaling receptors in microdissected TC and TIL. **A.** RNA-sequencing derived gene expression (log scale) of 12 T cells co-stimulation markers**. B.** RNA-sequencing derived gene expression (log scale) of 12 T cells co-inhibition markers.

### Validation by quantitative RT-PCR of selected genes

We used quantitative RT-PCR (qRT-PCR) to confirm gene expression of randomly selected genes among the most significantly upregulated in TC versus TIL and TIL vs TC. Ten genes were evaluated: five genes upregulated in TIL (*CD28, CCR7, CD79B, FCMR and PAX5*) and five genes upregulated in TC (*ELF3, MAL2, MUC1, TFAP2A and GPR37*). Overall, qRT-PCR (Figure 5B) results mirrored the RNA-sequencing data (Figure 5A).

**Figure 5:**
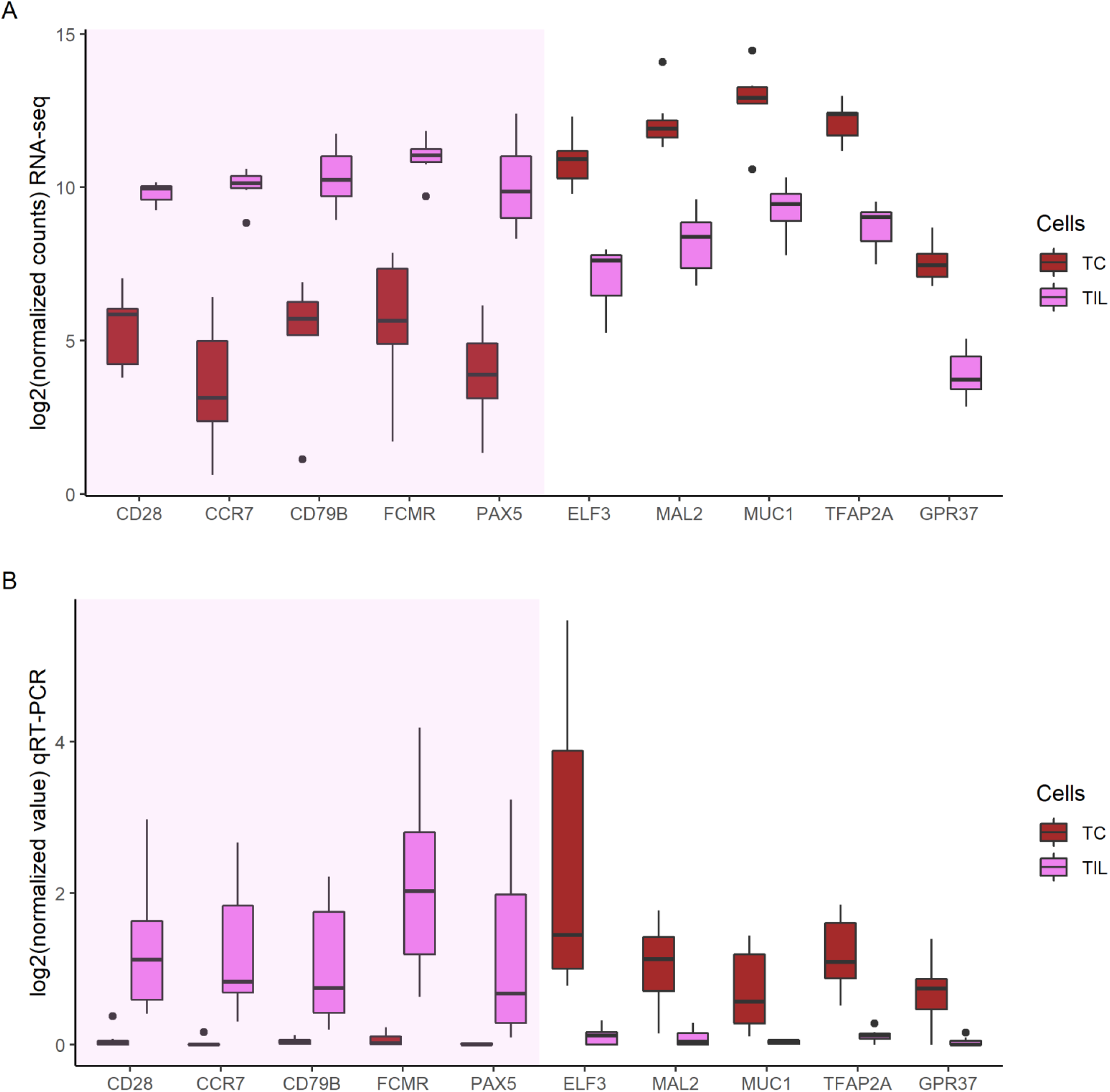
Validation of RNA-sequencing data by qRT-PCR. Expression level of ten randomly selected genes (5 upregulated in TIL and 5 upregulated in TC) were used to validate the RNA-sequencing data. **A.** RNA-sequencing derived gene expression (log scale) of 10 genes. **B.** Genes expression level by qRT-PCR (log scale) on 10 genes.

### Abundance of immune cell subpopulations

Deconvolution of bulk-RNAseq data was performed using microenvironment cell population (MCP)-counter method in order to estimate the abundance of different immune cell subpopulations (6). The analysis showed enrichment in adaptive immune cell signatures such as B cells and T cells in microdissected TIL, whereas innate immune cells signatures like neutrophils and monocytes were heteregenously present in TC and TIL (Figure 6A). Spatial enrichment in specific immune cell subpopulations was variable across the seven cases of TNBC. For instance, patient 73 (grey full arrows, Figure 6A) had TC enriched in T cells signature, particularly CD8^+^ T and cytotoxic T cells. This was confirmed by multispectral immunohistochemistry (IHC) that showed massive intraepithelial accumulation of CD8^+^ cells (Figure 6B). This case would be classified as “fully-inflamed” (20). On the opposite, patient 69 (black full arrow, Figure 6A) had low T cells and CD8^+^ T cells scores in TC but not in TIL. Multispectral IHC confirmed the absence of intraepithelial CD8^+^ cells (Figure 6B **and** Supplementary Figure 5A), that were rather present at tumor margins (Supplementary Figure 5B). Thus, both MCP-counter and IHC suggest that patient 69 has CD8^+^ T cells that are not located on the tumor core but rather at tumor margin, designated as “margin restricted”(20). High spatial variability was also observed in the monocytic lineage. Monocytic lineage score was high among microdissected TC in patient 65 (grey empty arrow, Figure 6A) and low in patient 88 (black empty arrow, Figure 6A). Multispectral IHC confirmed the accumulation of intraepithelial CD68^+^ cells in patient 65, whereas they were mainly in the stromal compartment surrounding TC in patient 88 (Figure 6B). Overall, these data validated the combination of LCM and MCP-counter deconvulation as an approach for spatial characterization of immune infiltrate in TME.

**Figure 6:**
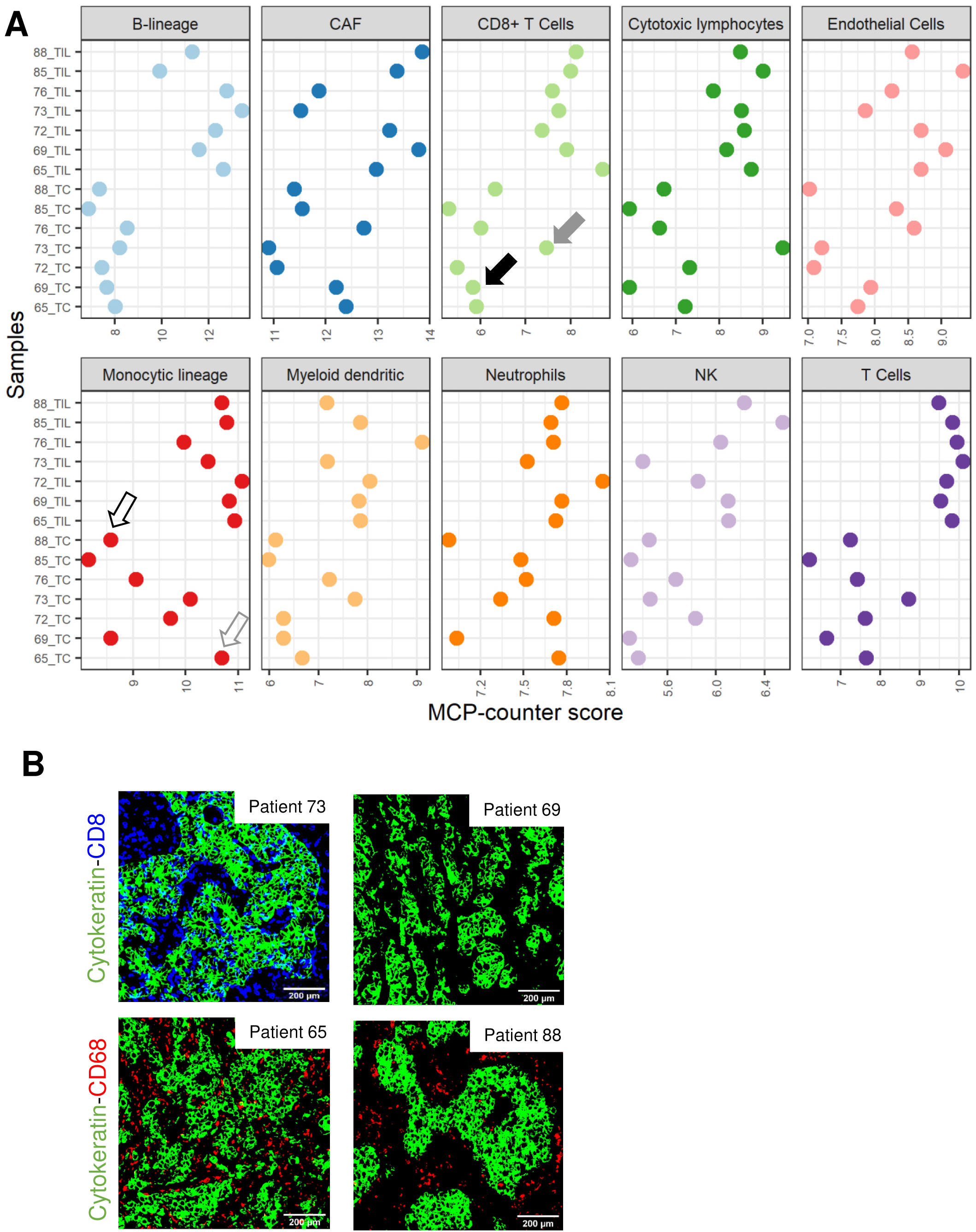
Abundance of immune cell subpopulations in TIL and TC estimated by MCP-counter. **A.** Dot-charts of eight immune cell subpopulations, cancer-associated fibroblasts and endothelial cells estimated by MCP-counter in paired microdissected TC and TIL (n=7). Grey empty arrow (patient 69) and black empty arrow (patient 88) indicate TC samples that are enriched or poor in monocytic cells, respectively. Grey full arrow (patient 73) and black full arrow (patient 69) indicate TC samples that are enriched or poor in CD8^+^ T cells, respectively. **B.** Representative multispectral IHC images showing the presence (patient 65) or absence (patient 88) of intraepithelial CD68^+^ monocytic cells. Multispectral IHC showed the presence (patient 73) or absence (patient 69) of intraepithelial CD8+ T cells. Epithelial cells are Cytokeratin^+^.

## Methods

### Samples and inclusion criteria

FFPE samples of TNBC were provided from the division of clinical pathology, Hôpitaux Universitaires de Genève. The research protocol was implemented in accordance with the declaration of Helsinki and Swiss law on biomedical research and was approved by the ethics committee of Geneva (CCER: 2018-02333).

### Histology

FFPE tissue sections were cut at 4 μm, floated onto a 45°C water bath, and mounted on Thermoscientific Superfrost® glass slides (Thermofischer, Massachusetts, USA). Tissue sections were dried overnight, dewaxed by immersion in xylene, rehydrated in ethanol of decreasing concentrations and stained with H&E. Sections were washed, dehydrated using increasing concentrations of ethanol and xylene and coated with a glass coverslip. Tissue sections were scanned using Pannoramic 250 Flash III scanner (3D Histech, Budapest, Hungary). All H&E sections were reviewed by a board-certified pathologist (J.C.T.).

### Estimation of number of cells per area

An H&E section of TNBC was scanned into MRXS image and quantified with QuPath (version 0.1.2, open source software). The polygon and point tools were used to count the nuclei within surface areas varying between 0.03-0.5 mm^2^. A linear regression of these values was used to estimate the number of cell for larger area of 1-2.5 mm^2^. Estimation of the quantity of RNA was calculated by multiplying the estimated number of cells to the estimated amount of RNA within a cell (10-30 pg/cell)(12).

### Workflow of FFPE laser capture microdissection and RNA-sequencing

To study spatial transcriptomics of TME on FFPE samples of TNBC, we performed LCM of 3 different cell subtypes (TC, TIL and Fib) followed by RNA extraction, library preparation and exome-capture RNA-sequencing (Figure 1). LCM procedure consisted in the manual delimitation of TC, stromal TIL (tumor core and margins) and Fib under direct microscopic visualization (step 1). Cells of interest were selected using a drawing tool on the Leica Laser Microdissection software. Selected regions of interest were cut from tumor sections with a laser beam. Microdissected samples were collected within the cap of a PCR tube placed under the polyethylene terephthalate membrane (PET) slide; the corresponding total RNA was extracted (step 2), followed by library preparation, which included the conversion of RNA into cDNA fragments and the addition of adaptors (step 3). Libraries were subjected to sequencing (step 4).

### Laser capture microdissection

Metal frame slides mounted with PET (Leica, Mannheim, Germany) were pretreated by coating with poly-lysine (Sigma-Aldrich, St-Louis, United States) and exposure to UV (15 mW/cm^2^/s) for 30 min. Consecutive sections from FFPE blocks were cut just before microdissection at 8 μm, floated onto a 45°C water bath and mounted on PET slides. After drying for 30 min, tissue sections were deparaffinized in xylene, rehydrated with decreasing concentrations of ethanol, stained with hematoxylin, dehydrated with increasing concentrations of ethanol, and cleared in xylene. A minimal surface of 2.5 mm^2^ of each cell population (TC, Fib and TIL) was microdissected using the Laser Microdissection Systems Leica LMD7000 and collected in separate 0.5 mL RNAse-free tubes. Laser parameters were set as follows: laser power of 39 mW, a wavelength of 349 nm, pulse frequency of 664 Hz and pulse energy of 120 μJ.

### RNA extraction, quantification and fragment size analysis

Microdissected cells were lysed by incubation with proteinase K at 56°C and stirring at 400 rpm for 3h. RNA was extracted in an RNAse-free environment using Qiagen® RNeasy FFPE kit (Qiagen, Hilden, Germany) according to the supplier’s protocol, treated with DNase I, eluted in 15 µl of RNAse-free water and stored at −80°C. Total RNA was quantified using Qubit™ RNA HS Assay Kit and Qubit® Fluorometer (Thermo Fisher Scientific, MA, USA). RNA fragment size was analyzed with the 2100 Bioanalyzer (Agilent, CA, USA) using Eukaryote total RNA Pico assay (Agilent, CA, USA). In case of insufficient RNA concentration, samples were concentrated using a Vacuum Concentrator Centrifuge (UNIVAPO 100 ECH, Uniequip, Germany) to reach the minimal volume recommended by the TruSeq RNA Exome® kit.

### Exome-capture RNA-sequencing

Multiplex paired-end exome-capture RNA-sequencing was performed on 16 microdissected samples obtained from 7 TNBC tumor cells: TC (n=7), TIL (=7) and fib (n=2). Libraries were prepared using TruSeq RNA Exome® kit (Illumina Inc, CA, USA) according to manufacturer’s protocols. The input quantities of RNA were adjusted to the DV_200_ values (Supplementary Table 2) as recommended in the TruSeq RNA Exome® reference guide and technical note (No. 470-2014-001): 40 ng if DV_200_ between 50%-70% and 55 ng if DV_200_ between 30%-50%. Each library was paired-end sequenced (2 × 100 bp) in duplicate in two different flow cell lanes (50 Million reads per library in total) using the TruSeq SBS Kit v3-HS on a HiSeq2000 platform.

### Data processing

Quality control was performed on raw sequencing data using the FastQC tool (21). Paired-end reads were aligned to the human reference transcriptome Homo_sapiens.GRCh38.cdna.all.fa.gz (release 94, obtained from ftp://ftp.ensembl.org/pub/release-94/fasta/homo_sapiens/cdna/) using Salmon v0.11.4 (22). Salmon uses a quasi-mapping approach for the fast quantification of transcripts expression. For mapping, the --gcBias flag was used, which accounts for fragment-level GC biases (under-representation of some sub-sequences of the transcriptome) present in RNA-seq data. After mapping, R Bioconductor (R version 3.6.2 and Bioconductor version 3.10) packages tximport (version 1.14.0) and DESeq2 (version 1.26.0) were used (23, 24) to summarize the transcript-level and gene-level estimates and for further down-stream analyses (25).

The dataset was filtered by discarding genes that had very low expression (gene abundance count < 10, across all samples). Variance stabilization transformation was performed using the VST function (26–28) implemented into the DESeq2 package. To visualize the sample to sample variation, PCA was performed on the top 500 most variable genes using the built-in R function *prcomp(),* and the principal components obtained were visualized using ggplot2 (29). For hierarchical clustering, the top 1000 variable genes were selected from the transformed data matrix and estimated the sample-to-sample Pearson correlation distances using the cor() function. Hierarchical clustering was performed with a complete linkage method using *hclust()* R function. A heatmap was plotted using the *Heatmap()* function from the ComplexHeatmap R package (30).

### Gene expression analysis

Differential expression analysis was performed using the DESeq function from the DEseq2 R package (23) on gene-level abundance estimates from Salmon. The DESeq function estimates size factors, dispersion values for each gene, fits a negative binomial model and performs hypothesis testing using the Wald test. DESeq2 also performs multiple test corrections on the p-values obtained from the Wald test using the Benjamini-Hochberg (BH) adjustment method (31). Adjusted p-value (FDR < 0.05) and log fold change (lfcThreshold = 0.58, which corresponds to a fold change of 1.5) were used to identify differentially expressed genes between tumor cells (TC) and TIL samples.

### Functional analysis

In order to characterize variations in biological pathways in TC and TIL, GSVA was performed (32). For this purpose, KEGG, Biocarta and Reactome gene sets from the C2 collection of the MSigDB v6.2 (c2BroadSets) were used (33, 34). GSVA analysis was performed on the variance stabilized expression value matrix of 1776 differentially expressed genes (adjusted p-value<0.05 and FoldChange ≥ 1.5, i.e. log_2_FC>0.58). Limma (35) was used for the generalized linear model fitting on the GSVA output in order to identify enriched pathways between TC and TIL. The Benjamini-Hochberg (BH) adjustment for FDR was applied. Significant pathways were identified with the adjusted p-value cutoff<0.05 and absolute log fold change cutoff for the enrichment score (log_2_FC > 0.58).

### MCP-counter analysis

MCP-counter method (6) was used to characterize immune cell subpopulations in microdissected samples. MCP-counter estimates the abundance of eight immune cell populations and two stromal cell populations in heterogeneous tissues from transcriptomic data. The gene expression matrix was provided as an input for the MCP-counter algorithm and was executed via the immunedeconv() R package, which provides unified access to several other deconvolution methods (36).

### Real-time quantitative PCR

Selected marker genes were validated using qRT-PCR. Sybr green assays were designed with Primer3 (version 4.1.0) with default parameters. Amplicon sequences were aligned against the human genome by BLAST to ensure the specificity of the primers for the genes of interest. The oligonucleotides sequences are listed in Supplementary Table 6. Oligonucleotide (Invitrogen, Thermofisher Scientific) sequences can be obtained upon request for the human TFRC and ALAS1 genes. cDNA was synthesized from 5 ng of total RNA using the PrimeScript Reverse Transcriptase (Takara by Clontech) and a mix of random hexamers - oligo dT primer, following supplier instructions. Specific cDNA was pre-amplified using the PreAmp Master Mix of Applied Biosystems (Thermofisher, Massachusetts, USA): cDNA samples were diluted 4x, primer pairs were pooled and used at 60 nM each, 14 PCR cycles (95°C 15 secondes - 60°C 4 minutes) were applied. PCR reactions (10 μl volume) contained diluted amplified cDNA (5.5x), 2 x Power up SYBR Green Master Mix (Applied Biosystems), 300 nM of forward and reverse primers. PCR was performed on an SDS 7900 HT instrument (Applied Biosystems) with the following parameters: 95°C for ten minutes and 40 cycles of 95°C for 15 secondes - 60°C for one minute. Each reaction was performed in triplicate on a 384-well plate. Raw Ct values were obtained with SDS 2.2.2. Selection of the best housekeeping genes, Normalisation Factors, and Normalized values was calculated according to the GeNorm method (37).

### Multispectral immunohistochemistry

FFPE tissue sections were cut at 4 μm, floated onto a 45°C water bath and mounted on positively charged Thermoscientific Superfrost ® plus glass slides (Thermofischer, Massachusetts, USA). Tissue sections were dried overnight, dewaxed by immersion in xylene, rehydrated in ethanol of decreasing concentrations, stained by Harris hematoxylin for 1 min, and coated by a glass coverslip with an aqueous mounting solution. Tissue sections were scanned into MRXS images using Pannoramic 250 Flash III scanner (3D Histech, Budapest, Hungary). Glass coverslips were removed by immersion in hot water, and slides were subjected to heat-mediated antigen retrieval in pH6 citrate buffer for 10 min. Endogenous peroxidases, non-specific proteins, endogenous biotins and avidins were blocked with corresponding blocking solutions from Dako®. The first primary antibody was applied to tissue sections, followed by a biotinylated-secondary antibody and a streptavidin-HRP complex, and revealed by AEC chromogen. Tissue sections were scanned and AEC staining was removed by immersion in ethanol of increasing concentrations. Antibodies were stripped by boiling tissue sections in pH6 citrate buffer for 10 to 20 min, and putative residual antibodies were blocked by Fab fragments directed against the host of the previous primary antibody used. Then, multispectral IHC consisted in sequential cycles of staining with primary antibodies reveled by AEC chromogen, tissue section scanning, removal of AEC chromogen with ethanol, antibody stripping and blocking with Fab fragments. Primary antibodies used in multispectral IHC were panCytokeratin (clone AE1/AE3, Dako®, dilution 1: 100), CD68 (clone M0876, Dako®, dilution 1: 100) and CD8 (clone, C8/144B, Dako®, dilution 1: 20).

### Image processing

MRXS scans were converted into TIFF files at half of the original resolution. A Matlab algorithm was written to execute all the following steps. Positive control tissues were cropped from the original slide image. For each tumor, images of CD8^+^, CD68^+^ and cytokeratin^+^ stainings were aligned to the image of hematoxylin staining. Region of interest were defined and processed to extract the positive staining (revealed by AEC chromogen) from each marker. CD8^+^ staining or CD68^+^ staining were superimposed to Cytokeratin^+^ staining for each tumor’s section in order to allow the visualization of final multispectral IHC images.

### Data availability

Upon publication, RNA-sequencing data will be available in raw FASTQ format from the Array Express database. The array express accession number is E-MTAB-8760.

## Discussion

We developed a new, robust method for spatial transcriptomic characterization of tumor microenvironment on FFPE samples. Unlike bulk RNA-sequencing on fresh tissue or single-cell RNA-sequencing, in which RNA is extracted without visualization of the tissue context, LCM focuses on specific regions or cells of interest. Here, we successfully applied this method to TNBC samples, the most aggressive subtype of breast cancer (38), characterized by the presence of TIL mainly in the stromal compartment with prognostic and predictive value (39, 40).

RNA is most often extracted from fresh frozen tissue. In most studies using laser microdissection, library preparation was performed on fresh frozen tissue (41, 42) and did not face the challenges of FFPE-related nucleic acid degradation. Fresh/frozen samples are not routinely available for all cancers and require long-term storage facilities with associated maintenance costs. Clinical pathology archives contain numerous FFPE tumor samples, collected from routine clinical practice. Such archives can date back to several years, even decades, making extraction of high-quality RNA challenging. Because neoadjuvant chemotherapy is the preferential treatment in TNBC (43), access to chemo-naïve samples is becoming scarce, limited to FFPE core needle biopsies. Our method is particularly relevant in this context. Beyond TNBC, our approach could be applied to investigate predictive biomarkers for response to immune checkpoint inhibitors administered either in neoadjuvant or metastatic setting, where access to tumor samples is limited to core needle biopsies (44, 45). The blocks we used were 3 to 10 years old and are representative of the material available in a clinical pathology biobank.

The most common protocol for RNA sequencing consists of enrichment of messenger RNA (mRNA) by poly(A) selection, which targets the polyadenylated RNA tail (46). However, poly(A) selection does not perform well when mRNA is degraded, because the poly(A) tail can be lost. The exome-capture protocol performs better than ribosomal RNA depletion on highly degraded and small amounts of RNA (47). Exome-capture on RNA extracted from FFPE showed minimal differences when compared with matched fresh-frozen samples (46, 48) and seemed the most appropriate approach for our method.

We applied spatial transcriptomics to either core needle or surgical biopsies of 7 FFPE samples of TNBC. PCA showed a clear separation between TIL, cancer cells and fibroblasts. This was further confirmed by per-sample GSVA analyses. Differential expression analyses showed that *MUC1* was among the most significantly upregulated gene in TC. MUC1 (CA15-3) is a well-characterized transmembrane mucin previously reported to be upregulated by 90% of TNBC (17) and it is FDA-approved serum tumor marker for monitoring metastatic breast cancer patients. Overxpression of *MUC1* validates the accuracy of gene-expression profile of microdissected TC. A similar approach could be used for identifying new biomarkers for diagnosis and prediction of response to therapy.

We questioned the abundance of different immune cell types from the expression profile with MCP-counter deconvolution (6), a well-established method that has been used to identify predictive biomarkers for response to immunotherapy (44). Spatial enrichment in specific immune cell subpopulations was variable across the seven cases of TNBC as confirmed by multispectral IHC. Stroma TIL were enriched in B and T cell signatures whereas myeloid and monocytic lineage signatures were heterogeneously enriched in TIL and TC. Here we propose that combining LCM and MCP-counter could be an alternative approach to multispectral IHC.

The repertoire of co-signalling receptors expressed on T cells is highly versatile and responsive to changes in the tissue microenvironment (18). An exploratory analysis of the spatial distribution of co-signaling receptors revealed that T cells co-inhibitory molecules/immune checkpoint inhibitors *BTLA* but not *CD160* was significantly upregulated in stromal TIL. Since BTLA and CD160 are both ligands of the same receptor HVEM (18), BTLA but not CD160, could facilitate immune evasion in the context of TNBC and hence represents a potential therapeutical target. Our approach of spatial characterization of co-signaling receptors through parallel analyses of the receptors and their ligands could bring new information on the cross-talk between components of TME.

Overall, our method allowed the concomitant analysis of: 1) the abundance of immune cell types; 2) the expression levels of co-signaling molecules in different TME compartments and 3) the enrichment of specific genes/pathways in cancer cells. Thes informations cannot be obtained by alternative solutions like nanostring® or multispectral IHC. It has potential applications beyond immuno-oncology and could be used in any FFPE tissue without predefining cell lineage-specific markers.

The method we developed has several limitations. Although fibroblasts represent a substantial proportion of the total surface of TNBC (5), we were able to collect sufficient RNA from microdissected fibroblasts in only 2 out of 7 TNBC cases. Fibroblasts are embedded within the extracellular matrix and are particularly difficult to isolate. A recent technical report on single-cell RNA sequencing showed that extended collagenase incubation time is required to release fibroblasts from tissue samples whereas non-adherent cells such as immune cells are readily and rapidly isolated by enzymatic disaggregation (49). It would be important to test whether adding digestion with collagenase could increase RNA yield of microdissected fibroblasts. Another limitation is intraepithelial TIL (patient 73 in our cohort) which are more difficult to microdissect than stromal TIL. To overcome this challenge, an alternative could be combining LCM with either single-cell RNA-sequencing (50) or IHC staining as we previously described for DNA whole-exome sequencing (9).

## Conclusion

In summary, we demonstrate the feasibility of using FFPE blocks of various ages to investigate the spatial transcriptomic profile of TME in TNBC. Optimization of LCM and RNA extraction protocols enabled the use of a small quantity of FFPE-derived RNA for exome-capture sequencing. The expression profiles of cancer cells, TIL and fibroblasts were clearly distinct, confirming the accuracy of microdissection. Spatial enrichment analysis revealed high variability in immune infiltrate across tumor samples. Since the majority of clinical samples are stored in FFPE, the current method could facilitate their reuse for research or clinical purposes and gives access to a large number of samples that would have been unfit for analysis with previous methods.

## Acknowledgements

We thank Dr Mylene Doquier and Mr Didier Chollet from the plateform of Genomics at the Faculty of Medicine, UNIGE, for their technical support.

## Supplementary Figures and Tables

**Supplementary Figure 1:**
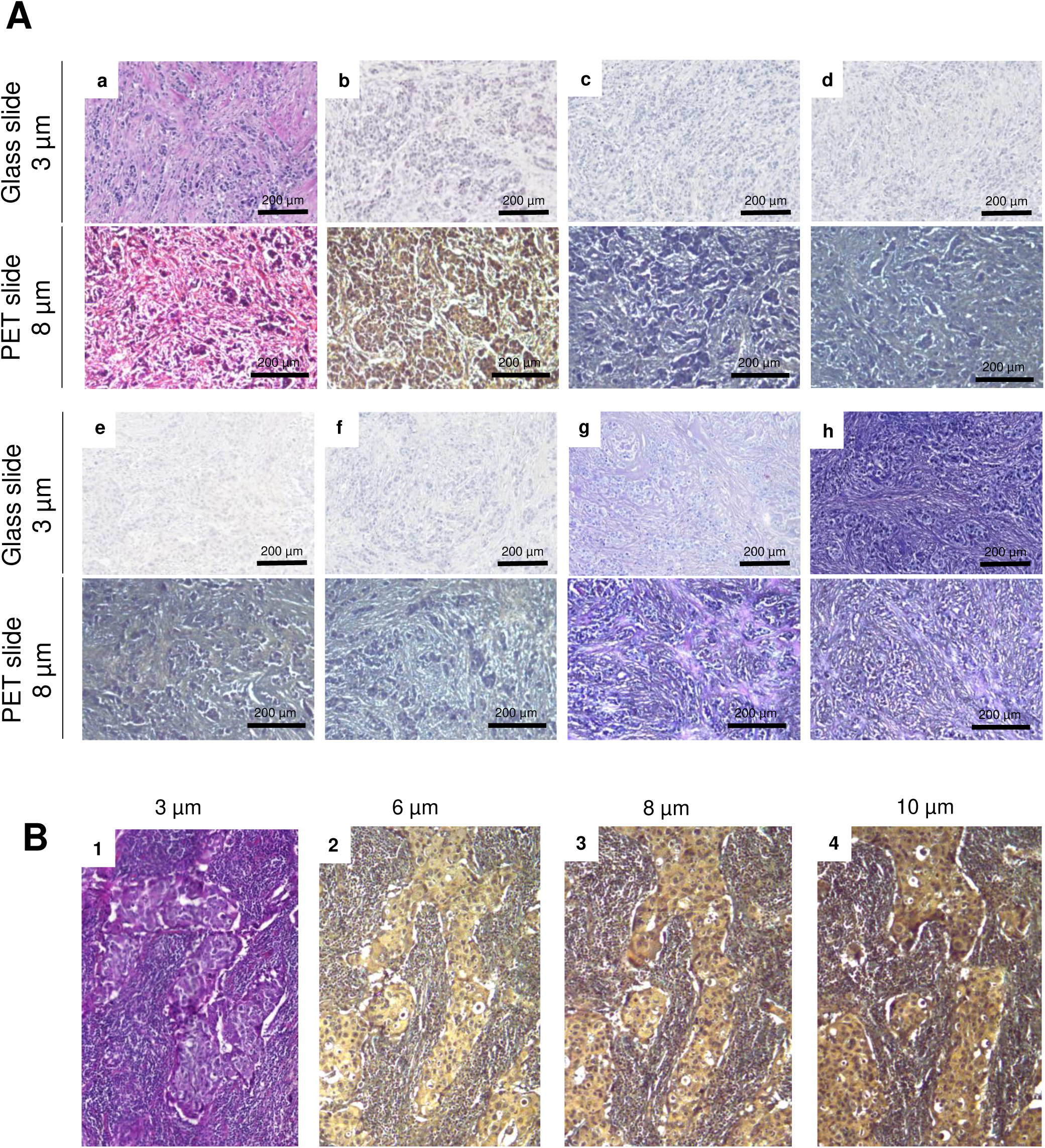
Optimization of staining protocol and tissue section thickness and staining before microdissection. **A.** Different staining protocols (a-h) performed on 3µm-thick tissue section mounted on glass slides (upper panel) and 8µm-thick tissue sections mounted on PET slides (lower panel). The optimal staining for subsequent LCM was obtained with protocol b. **B.** Tissue sections of different thickness (6-10 µm) stained with protocol b (panel 2-4) were mounted on PET slides and cell morphology was compared with the one obtained with H&E staining (panel 1).

**Supplementary Figure 2:**
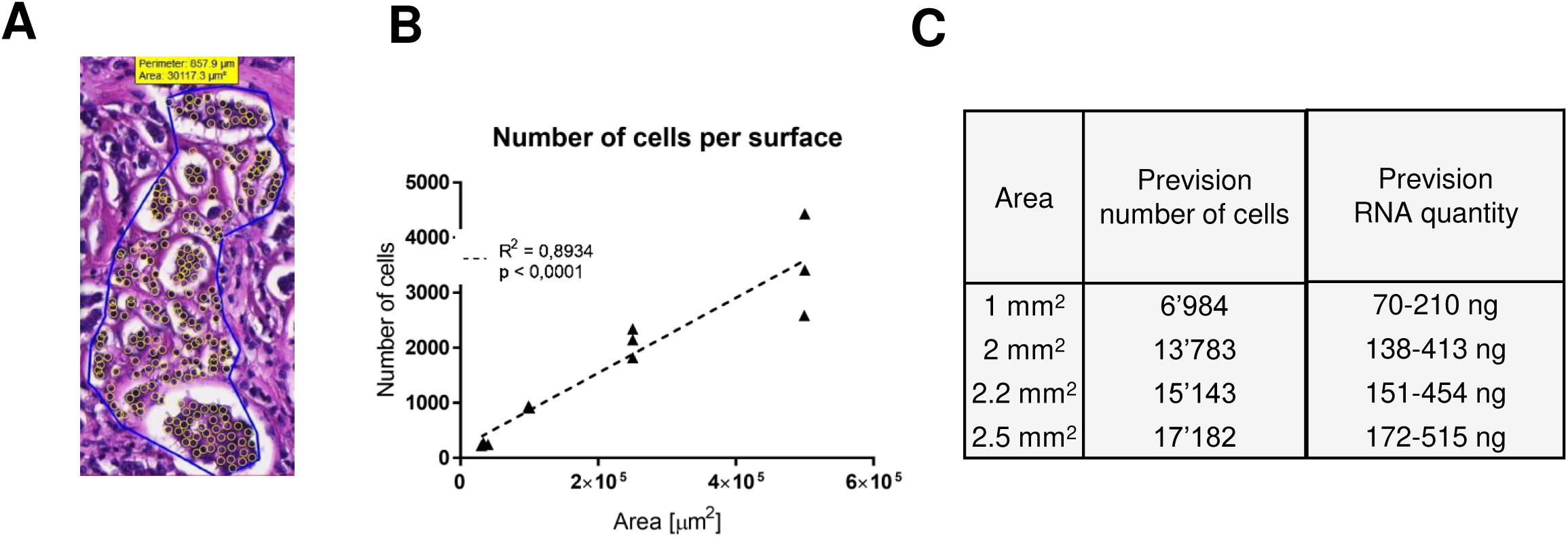
Estimation of RNA amount per area. **A.** Estimation of the number of nuclei on H&E section of TNBC with QuPath software. **B.** Linear regression curve showing correlation between the number of counted cells and surface area. **C.** Estimation of the expected amount of RNA according to the surface area.

**Supplementary Figure 3:**
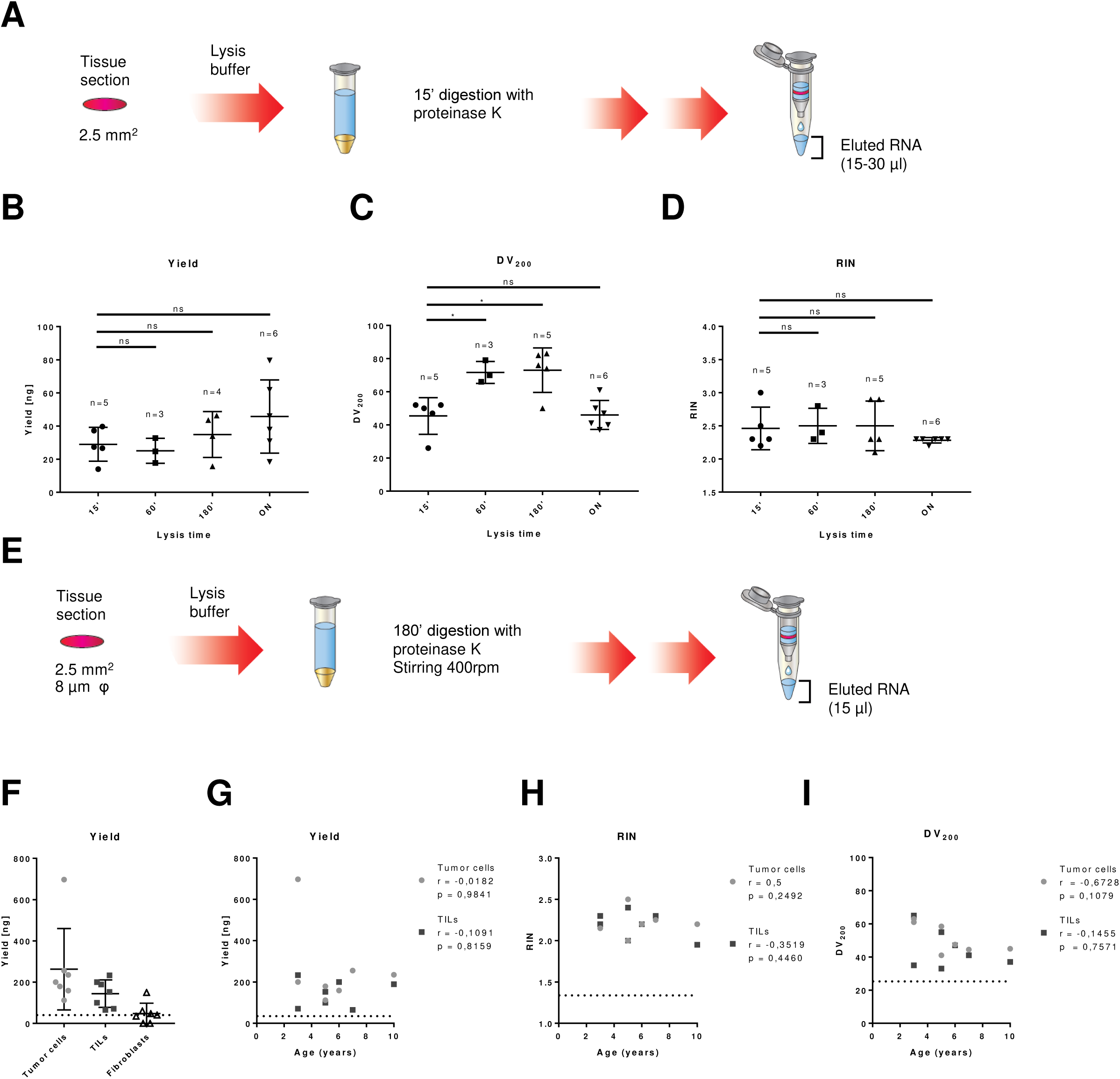
Assessment of RNA quantity and quality from microdissected TC and TIL. **A.** Original Qiagen RNeasy® FFPE Kit protocol**. B-D.**Yield, DV_200_ and RIN of RNA extracted with different durations of tissue digestion with proteinase K. **E.** Modified protocol for RNA extraction from the original Qiagen RNeasy® FFPE Kit protocol. **F.** Yield of RNA collected from microdissected TC, TIL and fib from 7 samples of TNBC. **G-I.** RNA yield, RIN and DV_200_ extracted from microdissected TIL and TC, according to the age of FFPE blocks. Dashed lines represent the minimal RNA quantity and quality required for subsequent RNA-sequencing. FFPE: formalin-fixed paraffin-embedded. RIN: RNA integrity number; DV200: percentage of RNA fragments >200 nucleotides.

**Supplementary Figure 4:**
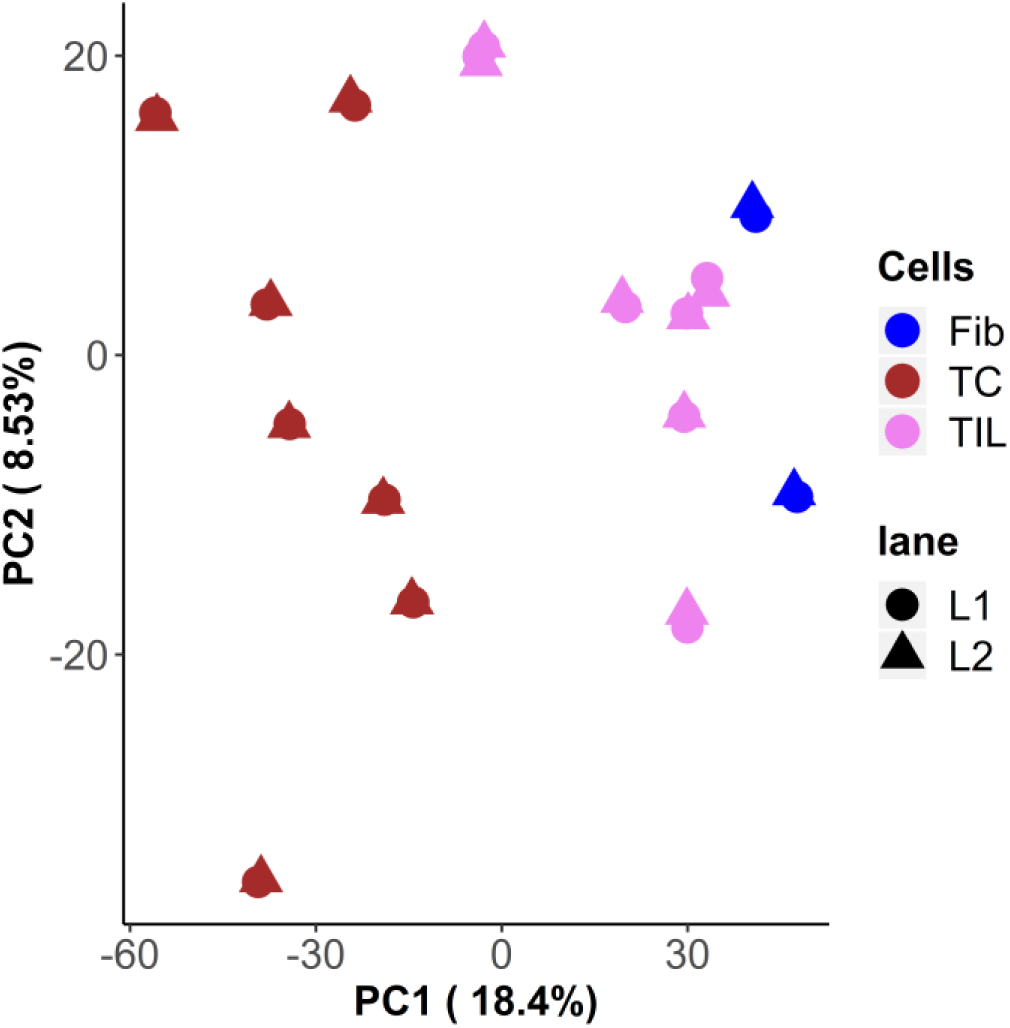
Technical replicates of principal component analysis (PCA). PCA of 16 microdissected samples representing the 500 most differentially expressed genes showing technical replicates (L1 and L2) from the multiplex mode of RNA-sequencing. TC = tumor cells, TILs = tumor-infiltrating lymphocytes, Fib = fibroblasts.

**Supplementary Figure 5:**
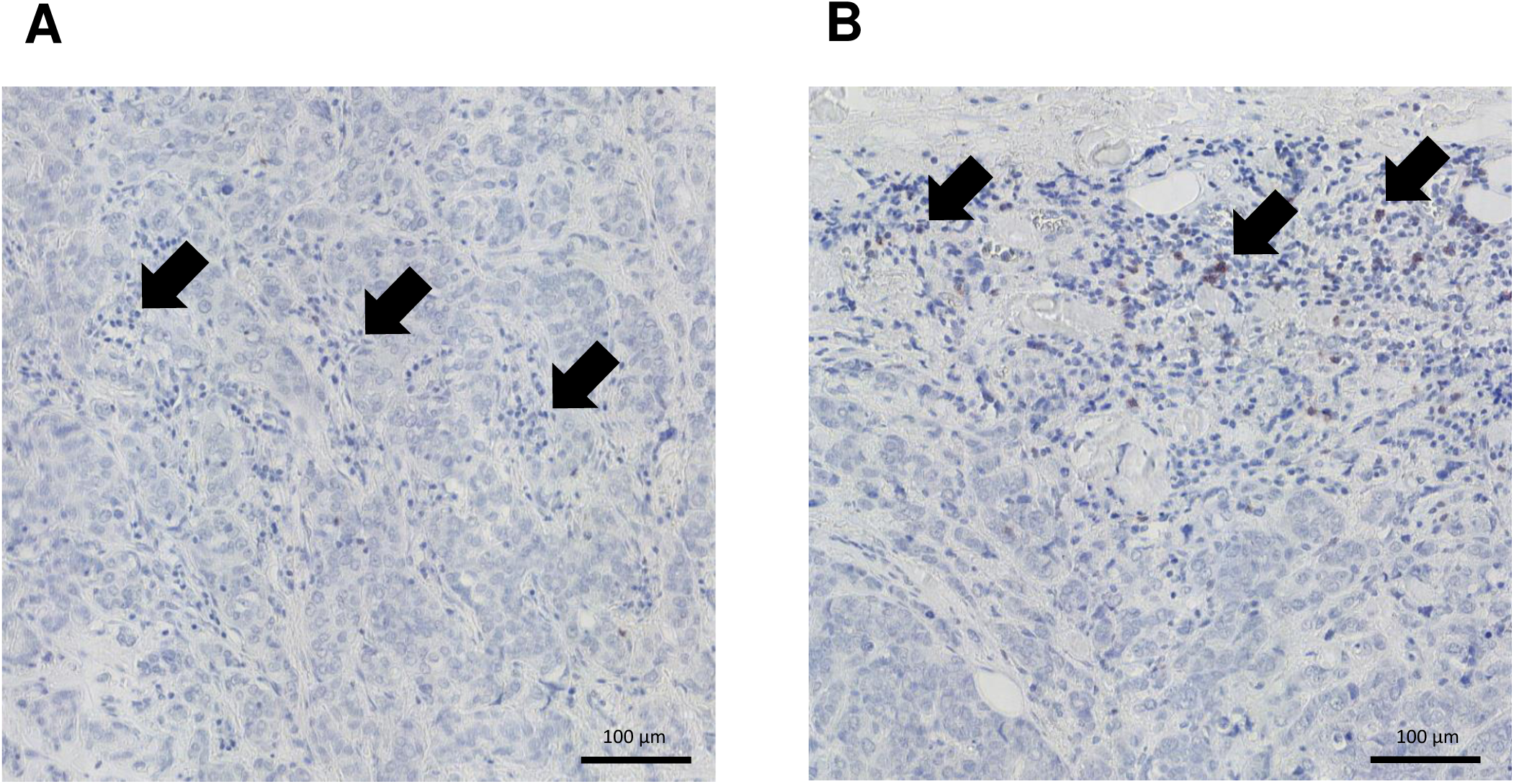
Immunohistochemical (IHC) staining of CD8^+^T cells in patient 69. **A.** CD8^+^ cells are rare in the tumor core. **B.** CD8^+^ TIL are present in the tumor margin. CD8^+^ TIL are pointed out with black arrows.

**Supplementary Table 1:** Tissue section staining protocols for LCM.

|  | dewaxing |  | rehydration |  |  |  | staining |  |  |  |  | washing |  |  | dehydration |  |  |  |  | Total time |
| --- | --- | --- | --- | --- | --- | --- | --- | --- | --- | --- | --- | --- | --- | --- | --- | --- | --- | --- | --- | --- |
| Protocol | Xylene | Xylene | 100% EtOH | 95% EtOH | 75% EtOH | H<sub>2</sub>O DEPC | Hematoxylin | Rinse and differentiate | Eosin | Cresyl violet 1% | Cresyl violet 4% | H<sub>2</sub>O DEPC rinse | H<sub>2</sub>O DEPC | Scott's tap water | 75% EtOH | 95% EtOH | 100% EtOH | 100% EtOH | Xylene |  |
| **a. Clinical pathology** | 10' |  | 10" | 10" | 10" | ☒ | 5' | 5' | 5' |  |  | ☒ | ☒ | ☒ | 10" | 10" | 10" | 10" | 10" | **27'** |
| **b. Labidi-Galy** | 1' | 1' | 1' | 1' | 1' | 2' | 2' | ☒ | ☒ | ☒ | ☒ | 3x | 1' | ☒ | 1' | 1' | 2' | ☒ | 1' | **15'** |
| **c. Labidi-Galy modified** | 1' | 1' | 1' | 1' | 1' | 2' | 2' | ☒ | ☒ | ☒ | ☒ | 3x | ☒ | 1' | 1' | 1' | 2' | ☒ | 1' | **15'** |
| **d. Espina** | 5' | 5' | 1' | 1' | 1' | 2' | 15" | ☒ | ☒ | ☒ | ☒ | 3x | ☒ | 1' | 1' | 1' | 1' | ☒ | 1' | **20'15"** |
| **e. Espina modified 1** | 1' | 1' | 1' | 1' | 1' | 2' | 15" | ☒ | ☒ | ☒ | ☒ | 3x | 1' | ☒ | 1' | 1' | 1' | ☒ | 1' | **12'15"** |
| **f. Espina modified 2** | 1' | 1' | 1' | 1' | 1' | 2' | 15" | ☒ | ☒ | ☒ | ☒ | 3x | ☒ | 1' | 1' | 1' | 1' | ☒ | 1' | **12'15"** |
| **g. Amini** | 5' | 5' | 30" | 30" | 30" | 10" | ☒ | ☒ | ☒ | 15" | ☒ | ☒ | 10" | 10" | 10" | 10" | 10" | 2' | ☒ | **14'45"** |
| **h. Clement-Ziza** | 5' | 5' | 30" | 30" | 30" | 10" | ☒ | ☒ | ☒ | ☒ | 15" | ☒ | 10" | 10" | 10" | 10" | 10" | 2' | ☒ | **14'45"** |

**Supplementary Table 2:** Quantity and quality of RNA obtained from microdissected cancer cells and TILs with optimized protocol for RNA extraction.

|  | Cell type | RIN | DV<sub>200</sub> | Yield [ng] | Quantity for RNAseq [ng] | Block age [years] |
| --- | --- | --- | --- | --- | --- | --- |
| Surgical biopsy 88 | Tumor cells | 2,15 | 63,00 | 200,20 | 40 | 3 |
|  | TILs | 2,30 | 65,00 | 233,10 | 40 |  |
|  | Fibroblasts | 1,97 | 55,33 | 47,04 | 40 |  |
| Surgical biopsy 73 | Tumor cells | 2,50 | 41,00 | 179,20 | 55 | 5 |
|  | TILs | 2,40 | 33,00 | 152,60 | 55 |  |
|  | Fibroblasts | N/A | 42,00 | 149,80 | 55 |  |
| Surgical biopsy 72 | Tumor cells | 2,2 | 47,5 | 158,9 | 40 | 6 |
|  | TILs | 2,2 | 47 | 200,2 | 55 |  |
|  | Fibroblasts | 2,3 | 37 | 60,2 | - |  |
| Surgical biopsy 69 | Tumor cells | 2,25 | 44,5 | 255,5 | 55 | 7 |
|  | TILs | 2,3 | 41 | 65 | 55 |  |
|  | Fibroblasts | 2,4 | 18,5 | N/A | - |  |
| Core biopsy 85 | Tumor cells | 2,15 | 61 | 697,2 | 40 | 3 |
|  | TILs | 2,2 | 35 | 71 | 55 |  |
|  | Fibroblasts | 2,2 | 45 | 35 | - |  |
| Surgical biopsy 65 | Tumor cells | 2,2 | 45 | 235,2 | 55 | 10 |
|  | TILs | 1,95 | 37 | 189 | 55 |  |
|  | Fibroblasts | N/A | 24 | 40,6 | - |  |
| Core biopsy 76 | Tumor cells | 2 | 58,5 | 112 | 40 | 5 |
|  | TILs | 2 | 55 | 100 | 40 |  |
|  | Fibroblasts | 2,1 | 62 | N/A | - |  |

**Supplementary Table 3:** FactQC summary. Total 64 FastQ files corresponding to the 16 samples sequenced in multiplexed (lane L1 and L2), paired-end (R1, R2 mode)

| QC parameter | nb<br>samples | nb<br>failure | nb pass | nb<br>warning |
| --- | --- | --- | --- | --- |
| Per base sequence quality | 64 | 0 | 64 | 0 |
| Per sequence quality scores | 64 | 0 | 64 | 0 |
| Per sequence GC content | 64 | 43 | 11 | 10 |
| Sequence Duplication Levels | 64 | 64 | 0 | 0 |
| Overrepresented sequences | 64 | 14 | 2 | 48 |
| Adapter Content | 64 | 4 | 48 | 12 |
| Per tile sequence quality | 64 | 32 | 0 | 32 |
| Per base sequence content | 64 | 32 | 0 | 32 |
| Per base N content | 64 | 0 | 64 | 0 |
| Sequence Length Distribution | 64 | 0 | 64 | 0 |

**Supplementary Table 4: List of genes upregulated in TIL**

**Supplementary Table 5: List of genes upregulated in TC**

**Supplementary Table 6:**
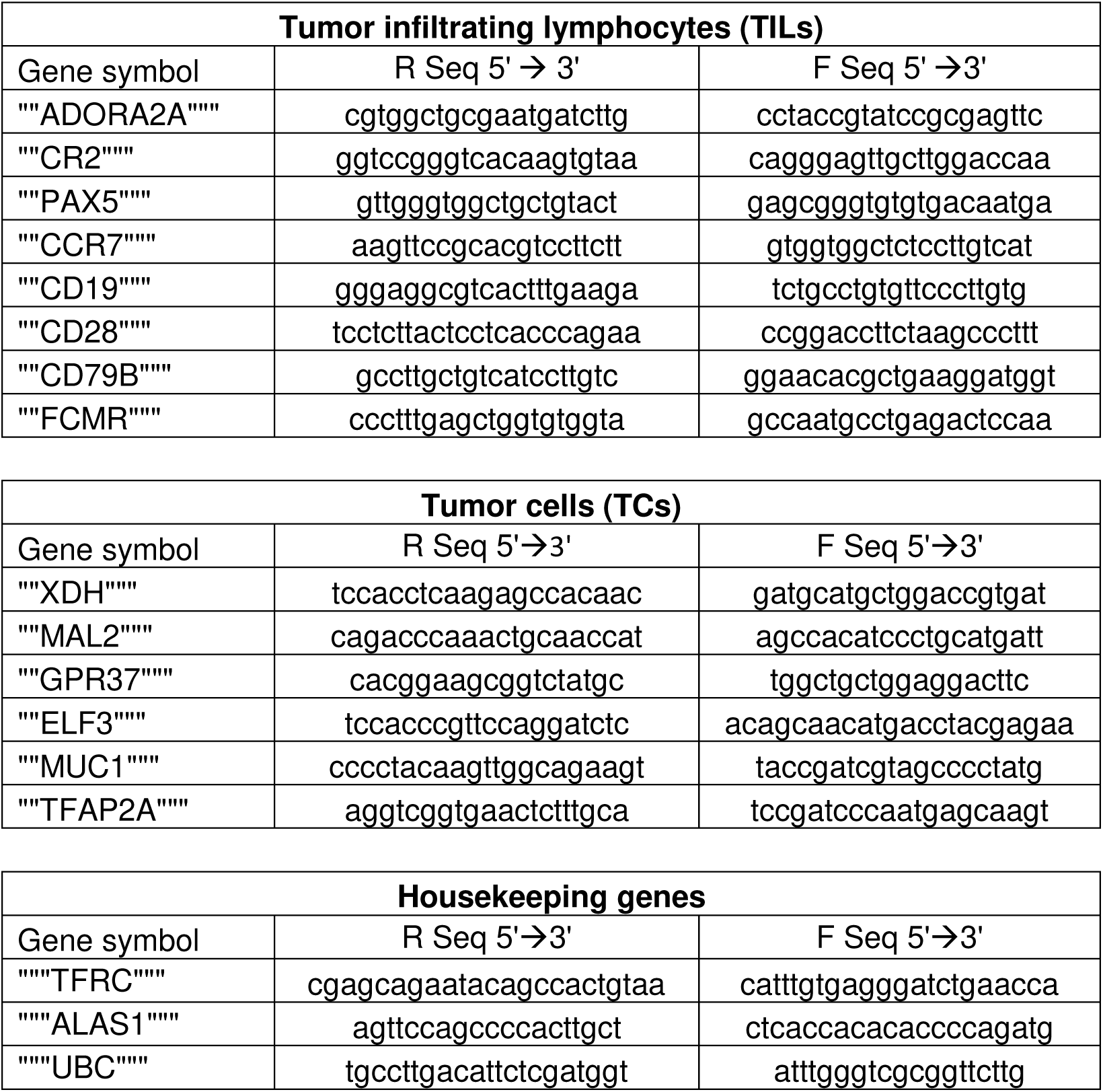
List of primers for qRT-PCR.

